# The physiology of deterrence: Flicker vertigo and its application in avian management

**DOI:** 10.1101/2024.06.20.599982

**Authors:** Takeshi Honda

**Affiliations:** Yamanashi Prefecture Agricultural Research Center, 1100, Shimoimai, Kai, Japan

**Author notes:** Correspondence: Takeshi Honda, Yamanashi Prefecture Agricultural Research Center, 1100, Shimoimai, Kai, Japan.

**Keywords:** agriculture, bird strike, conflict, Corvidae, flashing light, red

## Abstract

Human-bird conflicts are in a critical state, involving economic losses such as agricultural losses, bird strikes on aircraft and avian influenza. Traditional technologies leveraging bird vision and hearing often lose their effectiveness over time as birds become habituated to these stimuli. To address these challenges, our study introduces a novel countermeasure technology based on neurophysiology. The human brain reacts to flickering light, which can cause symptoms like headaches, nausea, and dizziness. In extremely rare cases, it can even lead to epilepsy. This led us to consider the possibility that similar stimuli could be applicable to birds. In our experiments conducted during the day, we used long-range flashlights. White flickering light had no effect on bird escape behavior. However, when cellophane film was attached to the flashlights to restrict the wavelength, the emitted red light induced escape behavior in birds. Additionally, employing two types of flashlights to generate flickering red+blue or red+green lights elicited escape behavior. However, the blue and green combination proved to be less effective. These results are highly similar to those found in human neurophysiology, showing that red light alone and the combination of red and blue lights have the most significant impact on the brain. By measuring the flight initiation distance (FID) of birds, we found that illuminated areas had a significantly higher FID (137m) compared to non-illuminated areas (12m). These findings suggest that applying principles of human physiology to wildlife management can offer new solutions for bird damage control.

## Introduction

Birds hold substantial significance owing to their ecological and cultural contributions; nevertheless, they present challenges in the realms of aviation safety, agriculture, zoonotic disease transmission, and the operational efficacy of wind energy installations (Conover 2001; Reidinger and Miller 2013). Additionally, birds can ignite wildfires by simultaneously contacting multiple power lines, resulting in their electrocution, subsequent heating, and combustion (Barnes et al. 2022). These wildfires can have a significant impact on forest ecosystems. One potential approach for mitigating these human-avian conflicts involves strategically dispersing birds away from high-risk human-centric environments. Documented strategies include the removal of birds from active aviation corridors (Blackwell et al. 2004, 2012), water bodies near poultry farms (Souris et al. 2014), agricultural lands vulnerable to crop depredation (Wang et al. 2019), and wind energy installations (Hodos 2003; Rebke et al. 2019). Although a range of technological methods such as lasers (Glahn et al. 2000; Elbers and Gonzales 2021), acoustic deterrence systems (Conover 1984), and bird-deterrent lines (Kessler et al. 1994; Honda 2012) have been employed, their efficacy remains limited. Notably, the effectiveness of some of these interventions appears to be species-specific; for example, lasers and bird-deterrent lines have demonstrated efficacy against the Anatidae family (Blackwell et al. 2002; Heim et al. 2022) and corvids (Honda 2012), yet their applicability is limited for other avian taxa. While acoustic deterrence has shown moderate success, it falls short in providing a long-term solution (Conover 1984; Gilsdorf et al. 2004). As such, the pursuit of universally effective measures to alleviate human-avian conflicts remains an ongoing challenge.

Given the widespread and effective use of electric fences in mitigating human-mammal conflicts (Tierson 1969; Honda 2022), the challenges of human-avian conflicts become evident. One might question why technologies aimed at reducing conflicts between birds and humans are ineffective. Electric fences, commonly used for mammalian deterrence, produce lasting aversion in animals that experience electrical shock, leading to long-term decreases in damage (Leblond et al. 2007; Reidy et al. 2008; Fischer et al. 2011). The electrical stimulus serves as an effective aversive cue because it is delivered each time an animal attempts an intrusion, thereby preventing habituation to the electric fence in mammals (Honda 2022). Electric fences are designed to complete an electrical circuit that involves the positive terminal of the electric energizer, the electric wire, the animal’s body, and the ground leading back to the negative terminal. While this design effectively delivers an electric shock to mammals, who are generally in contact with the ground, the same is not true for birds. Birds, especially when perched or in flight, are often not grounded, rendering electric shocks ineffective as a deterrent for them. Furthermore, beyond electrical stimuli, technologies employed for avian deterrence frequently induce habituation. Large noises generated by propane cannons (Conover 1984; Heim et al. 2022), as well as lasers that merely illuminate the vicinity of birds (Glahn et al. 2000; Brown and Brown 2021), act as neophobic stimuli but do not pose direct risks to birds. Consequently, birds become unresponsive to these stimuli over time (Blackwell et al. 2002). Escape behavior in birds is a trade-off between perceived risk and foraging; as the perceived risk diminishes—or in other words, as birds become unresponsive to stimuli—foraging becomes dominant, inhibiting escape behaviors (Lima et al. 1985; Verdolin 2006; Bonnot et al. 2017; Honda and Ueda 2023). As such, stimuli employed for reducing human-avian conflicts are not functioning adequately as aversive conditioning cues. This ineffectiveness likely contributes to the failure to significantly reduce conflicts.

Several types of conditioning exist, among which escape conditioning using aversive stimuli seems particularly suited to resolving human-avian conflicts. Traditionally, a diverse array of stimuli has been employed in conditioning. Electrical signals have long been used for this purpose (Mowrer and Lamoreaux 1946; Lovibond et al. 2009); however, conditioning stimuli are not confined to electrical impulses. For instance, flickering light is known to induce visual discomfort and flicker vertigo in humans (Walter and Walter 1949; Harding and Mills 1983; Rash 2004). Such flickering stimuli, especially at frequencies of 15-20 Hz, can cause headaches, dizziness, and nausea (Harding and Mills 1983; Wilkins 2016; Batra et al. 2019). Although it is extremely rare, affecting one in 100,000 individuals, these stimuli can induce photosensitive epilepsy in certain cases (Harding 1998; Da Silva and Leal 2017). This suggests that light stimuli exert influence not only on the eyes but also on the brain. Flickering stimuli are associated with large amplitude neural responses within the visual cortex (Gentile and Aguirre 2020); more precisely, the source of discomfort is transient abnormal synchronized activity of brain cells (Wilkins et al. 2010). Photosensitive epilepsy has been documented not only in humans but also in baboons, dogs, and poultry, indicating a lack of species specificity (Corcoran et al. 1979; Batini et al. 2004; Da Silva and Leal 2017; Szabó and Fischer 2021). Avians possess large eyes, the mass of which is determined by the size of the skull; specifically, avian eyes occupy the entirety of the skull, nearly touching at the midline (Brooke et al. 1999). Given these anatomical characteristics, visual deterrents may offer greater efficacy compared to deterrents targeting other sensory modalities. Indeed, strategies such as the deployment of lines with low visibility in agriculture and reducing visibility around feeding zones have been proposed for conflict mitigation (Honda 2012, 2015). Consequently, employing flickering light as an aversive stimulus in birds seems appropriate from both a physiological and wildlife management perspective.

While studies examining the effects of flickering stimuli on birds are scarce (Maddocks et al. 2001; Greenwood et al. 2004), extensive research has been conducted in humans. Both the frequency and the wavelength of flickering light have been studied, revealing that longer wavelengths (red) have a more profound impact on the brain compared to shorter wavelengths (blue) (Mesri and Dellepiane 1991; Takahashi and Tsukahara 1992). For the treatment of epilepsy patients, it is recommended to use blue sunglasses to attenuate the long-wavelength (red) stimulus (Takahashi and Tsukahara 1992). Furthermore, research has extended beyond single-wavelength stimuli to include the effects of two-color flickering light on the brain. It has been found that alternating flickering stimuli of red and blue induce a stronger neural response than red alone (Shirakawa et al. 2001; Parra et al. 2007). The strongest stimuli for humans are identified to be those that alternate between red and blue at 15 Hz (Parra et al. 2007). These studies have generally been conducted with light sources located close to the eyes (30-100 cm) and were aimed at minimizing health impacts on humans and other animals (Shirakawa et al. 2001; Parra et al. 2007). However, the use of high-power Light Emitting Diodes (LEDs) in spotlight-type, long-distance flashlights may enable the stimulation of birds even during daytime. Birds, similar to humans, possess blue and red-light receptors and can recognize both colors (Hart 2001; Peichl 2005; Tedore and Nilsson 2019); thus, the potential for applying flickering stimuli of red and blue is conceivable. Given these premises, the present study aims to verify whether escape behaviors can be induced in birds during daytime using various wavelengths of flashlight from a distance. Specifically, the study will focus on free-living corvids to: i) assess their responsiveness to long-wavelength light, and ii) evaluate the efficacy of alternating flickering lights of red and blue in inducing escape behaviors. The objective of this study is to provide an empirically substantiated approach to wildlife management that emphasizes strategies based on animal physiology.

## Methods

This study is composed of three experiments, each designed to both ascertain the effectiveness of flickering lights in aversive conditioning of avian species and to identify the most suitable wavelength for such stimuli. In humans, red light has been identified as the most potent monochromatic flickering stimulus, influencing not just the eyes but also the brain, thereby causing headaches and nonspecific complaints (Takahashi et al. 1981; Harding 1998; Parra et al. 2007). Furthermore, flickering lights alternating between red and blue have been demonstrated to elicit the strongest responses (Parra et al. 2007). Building upon these observations in humans, the study aims to examine the efficacy of light stimuli and avoidance reactions in corvids, specifically focusing on the Carrion Crow (*Corvus corone*) and the Large-billed Crow (*Corvus macrorhynchos*). All experiments were conducted during daylight hours.

### Experiment 1: Comparison of Monochromatic Light Stimuli

This initial study aimed to assess the aversive responses of crows to four types of flickering lights: white, blue, green, and red. A specialized long-range flashlight (Acebeam L18, Shenzhen Zenbon Technology Co., LTD, Shenzhen, China), which operates on a lithium-ion battery, was used for the study. The flashlight emits white light, produces a narrow beam, and can flicker at 8 Hz. To produce different colors, colored cellophane films (Komada Paper Co. Ltd, Ehime, Japan) were placed over the lens of the flashlight. This caused a reduction in the intensity of each color, with relative intensities quantified as 1 : 0.49 : 0.27 : 0.33 for white, blue, green, and red lights, respectively (Online source 1, upper panel). The experiment was conducted during daytime hours from January 20 to March 13, 2023. On each testing day, a random selection process determined the light type to be used, including a no-light condition, with the aim of collecting data from at least 30 individual subjects for each type of light. Surveys were conducted around agricultural test fields in Kai City and Minami Alps City, Yamanashi Prefecture, Japan. The selected survey route consistently measured 4.8 km in length. The site was located at an elevation of 280 m and included paddy fields and orchards near a river with a span of 359 m. To establish the flashlight as a first-order conditioned stimulus and the human operator as a second-order conditioned stimulus, researchers consistently wore orange vests.

The flashlight was positioned at a random distance between 20 and 200 m from the crows. After exposure, a laser rangefinder measured the exact distance (Nikon, Coolshot 20GII, Tokyo, Japan). Light exposure was restricted to 60 seconds. To minimize stress on the crows, the light was not used for tracking if they evaded. A video camera recorded behavior from the onset of light to evasion (HDR-CX370V, Sony, Tokyo, Japan). A higher deterrence effect from colored light compared to white light, despite its lower intensity, would indicate that color is the critical factor. Previous research indicates that avian sensitivity remains relatively consistent between 330-630 nm and sharply decreases above 630 nm (Hart, 2001). The red light used had a peak wavelength below 630 nm, suggesting that almost all tested colors were within the range visible to crows (Online source 1, upper panel).

### Experiment 2: Comparison of Red and Green Lights

Building on observations from monochromatic light experiments that showed green and red lights as aversive stimuli (see Results section), specialized flashlights (Archer M Pro Green or Archer M Pro Red, Maxtoch Co., Ltd, Zhejiang, China) were employed. These flashlights were also powered by lithium-ion batteries. Comparisons were conducted under three distinct conditions: monochromatic green light at a wavelength of 514 nm, monochromatic red light at a wavelength of 624 nm, and simultaneous flickering of both green and red lights. Similar to the flashlight used in Experiment 1, the devices in this experiment were also engineered for long-range illumination. In contrast to Experiment 1, the flickering frequency was not fixed but varied between 6 and 12 Hz, involving both high-frequency and low-frequency cycles. The use of a variable-frequency flashlight was due to the unknown optimal flickering frequencies for avian subjects. The flickering of the green and red lights was not synchronized; both lights could be either simultaneously illuminated or extinguished. Survey locations and methodologies remained consistent with those detailed in Experiment 1. Data collection was performed between March 14 and April 28, 2023. On each day of investigation, the type of light emitted was determined randomly, and six days of data were collected for each light condition. The wavelengths of color LED flashlights are shown in Online source 1 (lower panel).

### Experiment 3: Combining Multiple Colors in Flickering Lights

Building on human studies that indicate flickering lights alternating between red and blue elicit stronger responses than red light alone (Parra et al. 2007), this experiment presented crows with three different light conditions: blue+green, red+green, and red+blue. The study period ran from May 1 to July 4, 2023, using methods consistent with Experiments 1 and 2. Each light condition was maintained for a week and tested over nine days. For example, if three surveys were conducted in the first week, all were uniformly executed under the blue+green condition, followed by the red+blue condition in the next week. The order of these light conditions was randomized. The reason for maintaining uniform light conditions for a week was to account for the possibility of birds initiating evasion upon exposure to a specific type of light. For example, both blue+green and red+blue conditions contain blue light, and birds occasionally initiated evasion upon exposure to blue light. Keeping light conditions consistent within a week aimed to maximize the distinction between light types. The relative light intensities of the blue, green, and red monochromatic flashlights were quantified as 2.6 : 2.9 : 1, respectively (Online source 1, lower panel).

After completing the previous experiments, the flight initiation distance (FID) of the birds was measured on July 5-6. Researchers walked along the survey route and measured the distance at which crows began to evade using a laser rangefinder. To reflect the impact of second-order conditioning, orange vests were worn, similar to the light exposure experiments. Constant eye contact was maintained with the crows while approaching them on foot. This was because avian species respond to the direction of human faces (gazing), and the FID varies depending on whether the approaching human is facing them (Clucas et al. 2013; Yu et al. 2023).

Additionally, to measure the FID of crows unaffected by the light exposure experiments, surveys were conducted at locations 4-8 km away from the established survey sites. The researchers traveled by bicycle while searching for crows with binoculars and approached them on foot from approximately 300 meters to measure the FID.

### Statistical Analysis

The primary aim of this study is to investigate how different types of light affect avian evasion behavior. To address this objective, we included ‘type of light’ as an explanatory variable in our models. Additionally, we considered the ‘distance from the birds to the light source’ as another explanatory variable to account for the potential weakening of evasion behavior with increased distance from the light source. To quantify evasion behavior, the elapsed time from the moment of light exposure to the birds’ evasion serves as the dependent variable. This variable is right-censored at 60 seconds, indicating cases where birds remained exposed to the light for the entire observation period without evading. Given the censored nature of the data, traditional linear models are inadequate, thus necessitating the application of survival analysis techniques. Specifically, survival regression models were fitted using the ‘survreg’ function in R software (version 4.1.2). A log-normal distribution was assumed for the dependent variable to account for the right-censored nature of the data. To avoid the increased risk of Type II errors (false negatives) associated with corrections for multiple comparisons, we employed an exhaustive AIC-based model selection approach for grouping the light variables. This involved computing the AIC for all possible combinations of light variables and selecting the model with the minimum AIC value. To identify the optimal grouping for the five light color variables (e.g., experiment 1, Red, Blue, Green, White, and Control), a comprehensive combinatorial approach was employed. Variables were partitioned into multiple groups, and a separate survival regression model was fitted for each unique combination. The primary objective was to identify the combination that minimized the AIC value, providing the most statistically robust model.

For the comparison of FID, Welch’s t-test was employed.

## Results

### Experiment 1

No significant differences were observed between the white and blue flickering lights and the no-light condition (Fig. 1), as evidenced by survival analysis (white: p=0.99, blue: p=0.65). On the other hand, both red and green lights showed significant differences when compared to the no-light condition (p<0.03). As the distance from the light to the crow increased, the effectiveness of the light decreased (p=0.035). According to Akaike Information Criterion (AIC)-based analyses, red and green lights were categorized into distinct clusters separate from the no-light condition. Given these minimal p-values and the AIC results, it is inferred that the effects of red and green lights diverge from those of the no-light condition. In the current experiment, the intensity of colored lights was reduced by overlaying filters on the white light (light intensity: white > blue > red > green; refer to the Methods section). Nevertheless, despite their lower intensities, red and green lights were effective in eliciting aversive behaviors among crows. These results imply that the wavelength of the light could have a more significant impact on crow evasion behavior than the light intensity itself.

**Fig. 1.**
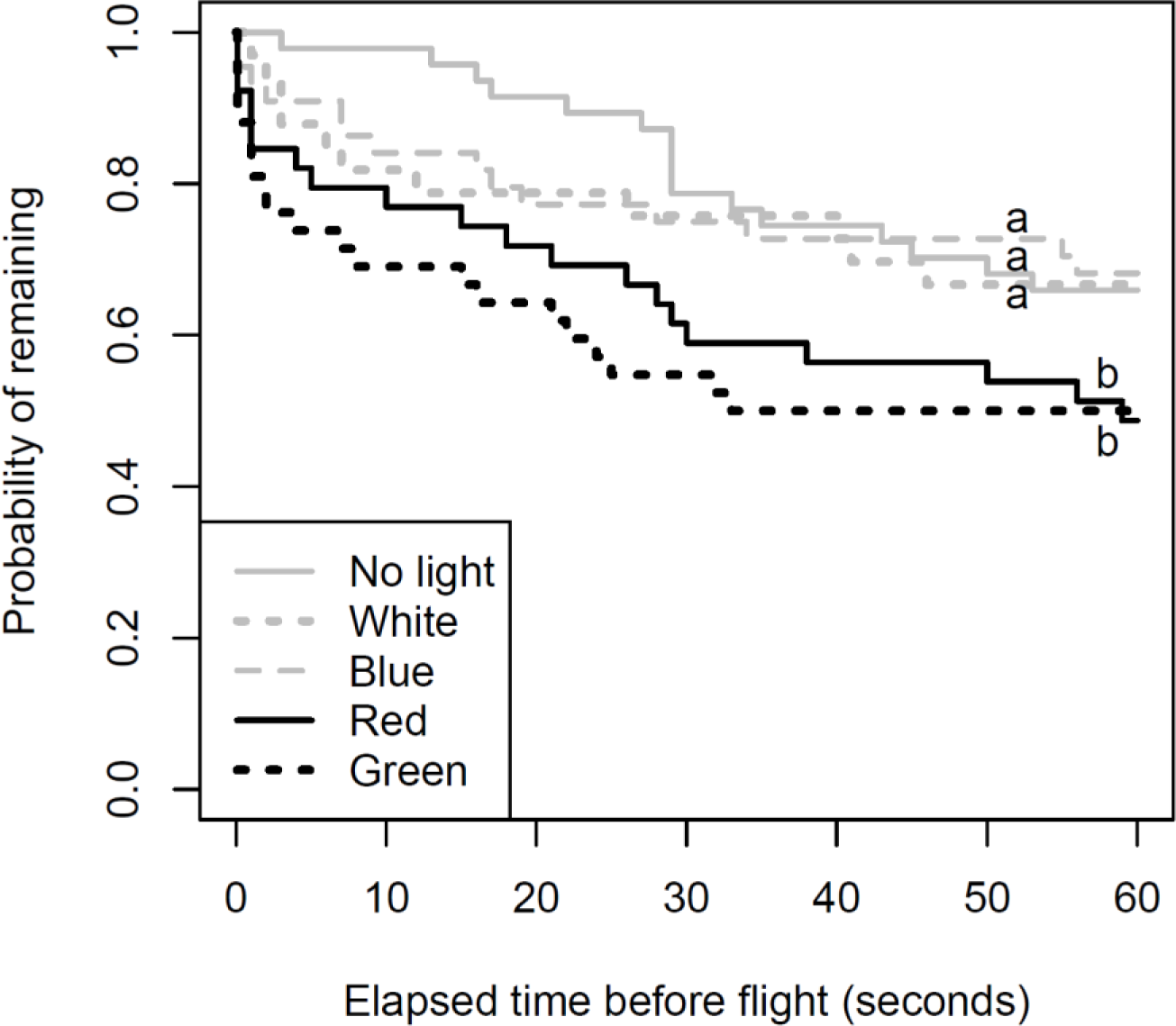
Variation in remaining probability for crows exposed to white LED strobe light (8 Hz) and colored filters. A flashlight emitting white strobe light was outfitted with blue, red, and green color cellophane filters and directed at the birds. The light conditions that demonstrated no difference from the control (no light) are represented with gray lines (p>0.05), while those that did demonstrate a significant difference are represented with black lines. Due to the use of color cellophane filters, the intensity of the white strobe light was the highest, whereas intensities were lower for the blue, red, and green lights. Alphabetic labels denote distinct treatment groups. It should be noted that the ‘probability of remaining’ under the control (no light) condition may vary over time, as crows may evade when observed by humans

In all experiments, including Experiments 1-3, crows showed no abnormal behaviour other than escaping.

### Experiment 2

In this experiment, color filters were not utilized; instead, LEDs emitting light at specific wavelengths (red and green) were employed. The number of observed crows was 91, 67, 77, and 64 for red, green, both colors, and no illumination, respectively. The flickering of the green and red lights was not synchronized; both lights could be either simultaneously illuminated or extinguished. These asynchronous flickering red and green lights exerted a strong influence on crows compared to the untreated condition (Fig. 2), as evidenced by survival analysis (p<0.001). Although the light intensity was greater for green than for red, the impact on crow aversion behavior was greater for red than for green. This inversion phenomenon suggests that light intensity alone cannot account for crow evasion behavior. Light stimuli incorporating both red and green had a comparable impact on crows as did the red light alone. Although no difference between red and green was observed in Experiment 1, the discrepancy does not arise as different types of lights were used in experiment 1.

**Fig. 2.**
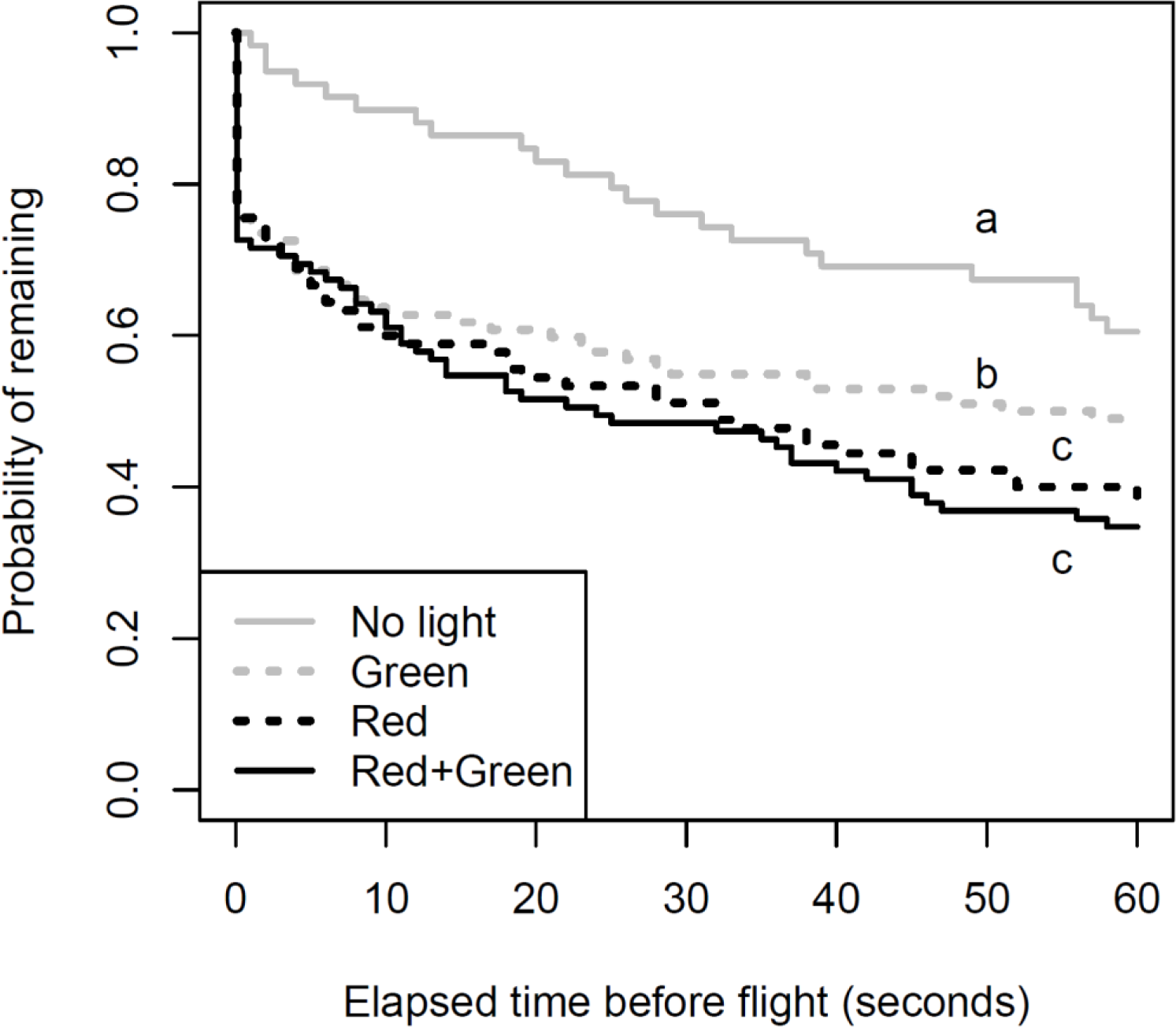
Persistence probability differences in crows exposed to variable-frequency strobe lights emitted directly from red and green LEDs, without the use of color filters. Lights operated between 6 Hz and 12 Hz and were not synchronized. Alphabetic labels denote statistically distinct treatment groups

### Experiment 3

Illumination of any light type increased aversion behavior in crows compared to the untreated condition (Fig. 3), as shown by survival analysis (p<0.005). The most effective combinations of light for causing aversion were red+green and red+blue, as determined by AIC-based model selection. The combination of blue+green had less effect on aversive behavior. When the flight initiation distance (FID) was compared, the crows in areas exposed to light showed a significantly higher FID than those in areas not affected by light (Fig. 4, t=5.78, df=13.3, p<0.001). In untreated areas, investigators could often approach directly below the crows perched on power lines, whereas the median FID in treated areas was approximately 137 meters.

**Fig. 3.**
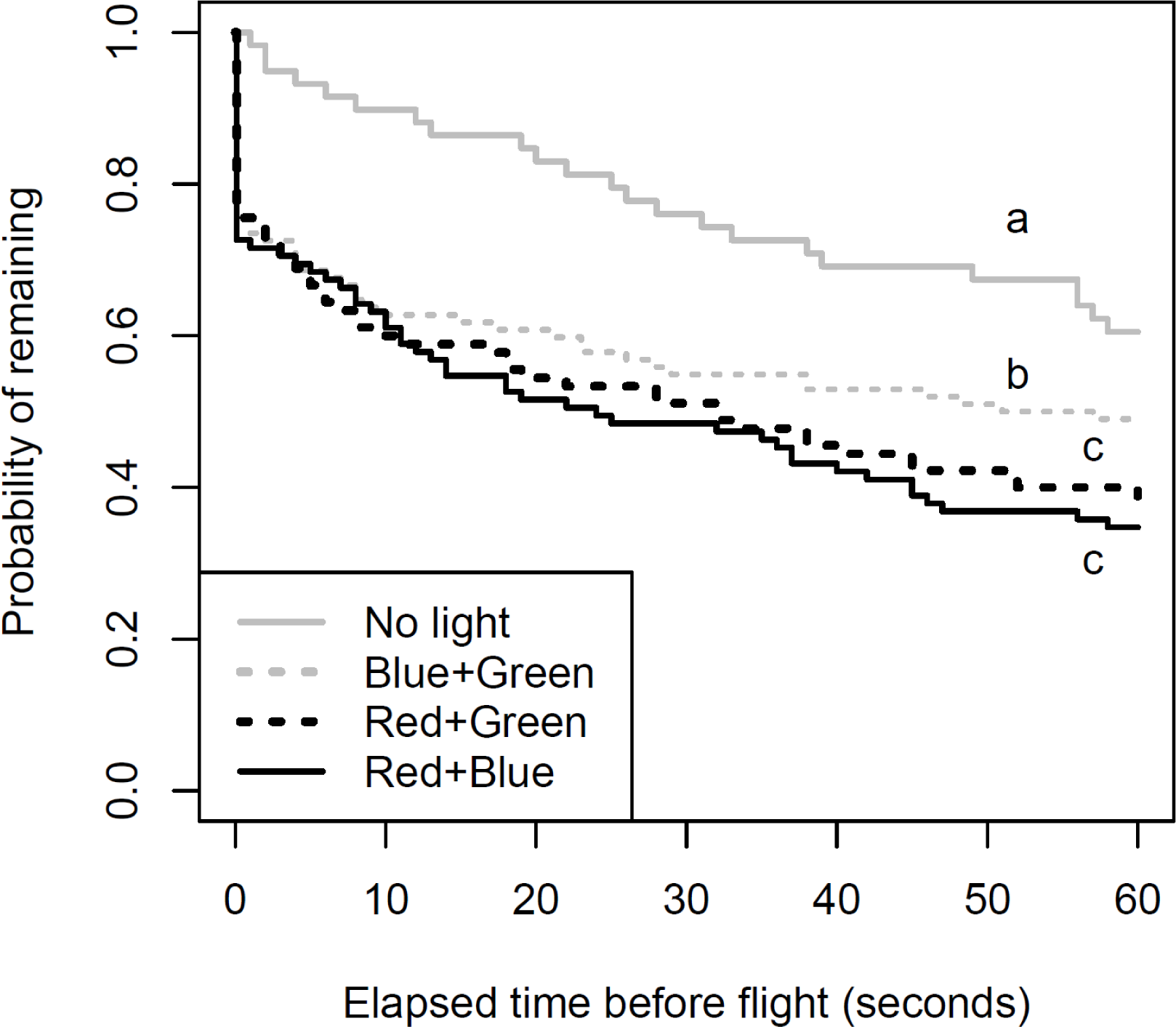
Variation in remaining probability for crows exposed to colored strobe light. The same flashlight as used in Fig. 2 was employed, with an additional blue LED flashlight incorporated into this experiment. Alphabetic labels denote statistically distinct treatment groups.

**Fig. 4.**
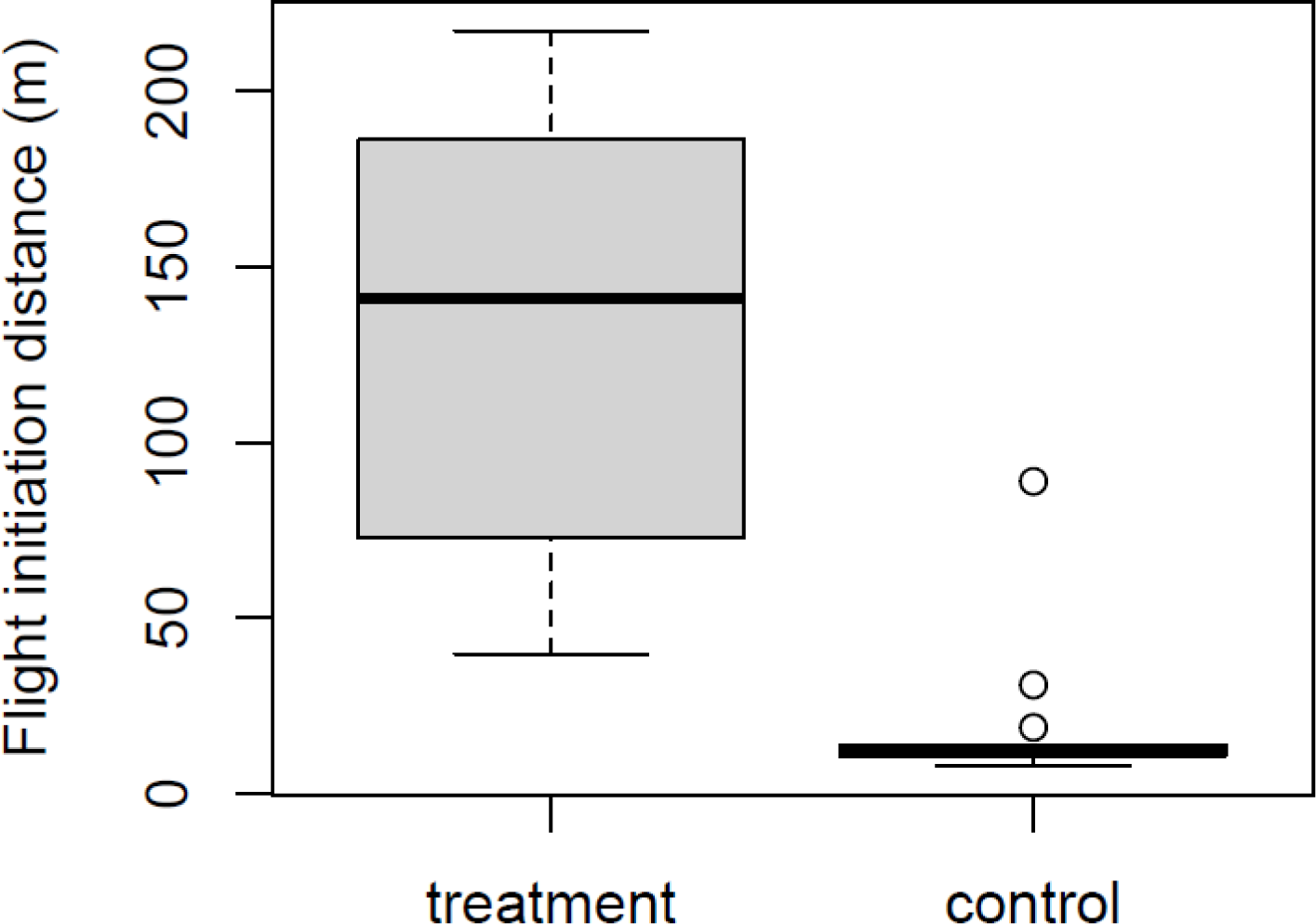
Comparison of flight initiation distance (FID) in crows between treated and untreated (control) areas. The boxplot illustrates the distribution of FID in crows from two distinct areas: one subjected to repeated strobe light exposure using a flashlight and another where no treatment (control) was applied. The investigation of FID was conducted immediately after all light exposure experiments were completed

The results were consistent with those from Experiments 1 and 2, confirming that the blue+green combination was less effective than combinations including red. Taking into account all the experiments, we can conclude that the single most effective light color for affecting crow behavior was red, and the most effective two-color combinations were red+green and red+blue. No clear difference was seen between the effectiveness of single colors and two-color combinations.

## Discussion

This study builds on previous research that has investigated the effects of flickering lights on human brain activity (Mesri and Dellepiane 1991; Tobimatsu et al. 1999; Parra et al. 2007). We extend this work by examining how various colors of light—white, blue, green, and red—affect birds. Results show that red light, which has a longer wavelength, strongly stimulates brain activity in birds. We also found that a flickering pattern of red and another color was more effective than a blue and green combination. However, we did not find difference between red-only and red + blue light in birds, suggesting that there might be subtle differences in how birds and humans respond to these stimuli.

To understand how two-color light patterns affect both birds and humans, it’s important to first know how light generally affects the brain. White light is a mix of different colors that include blue, green, and red light. Red light strongly influences brain activity because it specifically activates cells in the eye sensitive to red, which then send signals to the brain (Mesri and Dellepiane 1991; Harding 1998). Generally, stimulating single types of these eye cells leads to a stronger impact on the brain. In humans, there are three main types of these cells: one sensitive to blue light, another to green, and a third to red (Peichl 2005). Stimulating only the red-sensitive cells has a strong impact on the brain, although the reasons for this are not fully understood. While the sensitivity profiles of human cone cells for green and red overlap significantly (Harding 1998; Peichl 2005), the sensors for red and blue do not overlap, allowing for distinct detection. Therefore, alternating between red and blue light produces a more robust neural response, compared to red light alone (Shirakawa et al. 2001; Parra et al. 2007). This offers an explanation for why a combination of red and blue light is generally considered more effective than red light alone for stimulating brain activity in humans.

So why did our study find no difference between red-only and red + blue light in birds? One idea is that the blue light might lessen the effect of the red light. Earlier research shows that using red and blue lights one after the other has a strong effect on the brain, while using them together at the same time lessens this effect (Shirakawa et al. 2001; Parra et al. 2007). In our tests, we used two different light sources that were not synchronized. While past studies in humans have used alternating red and blue lights to study epilepsy, none have used the same kind of light stimuli as in our study. The lack of synchronization in our light sources could be why we found no difference. In our study, we lacked information on the most effective flicker frequency for stimulating birds. Consequently, the flicker was set to a variable frequency, which is why the two colors did not synchronize. Future work, with more precise control of light stimuli, will likely elucidate the effectiveness of using dual-color light.

Despite the subtle differences observed between humans and birds, it is intriguing that stimuli similar to those that stimulate the human brain also strongly stimulated birds. Human and avian eyeballs have different structures, with humans possessing three types of photoreceptors that can recognize blue, green, and red, while birds have four types, including the ability to recognize ultraviolet. That both species are strongly stimulated by red light, despite these differences, suggests that longer wavelengths universally provide stronger stimulation, irrespective of the types of photoreceptors present. Furthermore, there are differences in the brain regions responsible for processing vision between humans and birds. While humans process visual stimuli in the cerebrum, birds utilize the midbrain. Despite processing visual stimuli in different parts of the brain, it is intriguing that both species are strongly stimulated by long-wavelength light (red). In the case of photosensitive epilepsy in humans, it is not just the visual cortex in the cerebrum that is stimulated, but the stimulation propagates throughout the brain (Tobimatsu et al. 1999). It is possible that birds avoided the long-wavelength blinking light because the red light stimulated the midbrain and then propagated throughout the brain. If flicker vertigo is considered a very mild symptom of photosensitive epilepsy, the characteristic of blinking lights to stimulate the entire brain—unaffected by major structural differences between mammals and birds—may have broad applicability in wildlife management. It is known that Fayoumi chickens are susceptible to flickering light; however, to our knowledge, there is no research on the effects of light wavelength on the brain in avian species. The relationship between flickering light and light wavelength revealed in this study is beneficial for both wildlife management and physiology

To utilize the results obtained in this study for the effective technology in mitigating human-avian conflict, it may be essential to investigate the flickering frequency of light in greater detail. High-frequency flickering light may appear as if it’s a constant, steady light to the human eye. The specific frequency at which this flickering is no longer distinguishable and appears as constant light is known as the critical flicker frequency (CFF) (Rash 2004). Generally, the CFF in birds is higher than in humans, allowing birds to recognize higher frequency flickering light as flickering (Bateson and Feenders 2010; Boström et al. 2016). Separate cells in the eye transmit signals for the lighting (ON) and extinguishing (OFF) of light, and during flickering, ON and OFF signals are sent alternately. When this frequency becomes too high, the signals cancel each other out, rendering the flickering unrecognizable (Sharpe and Stockman 1999). The difference in CFF between humans and birds suggests that the flickering frequency that most strongly stimulates the brain may differ, warranting further investigation into effective flickering frequencies for conflict mitigation.

To achieve results similar to this study, it is possible to utilize commercially available flashlights (Maxtoch M Pro). There is potential to repel birds using these flashlights in various locations such as areas surrounding airports (bird strike), ponds near poultry farming facilities (avian influenza), places where birds inflict damage on grains and fruits (crop damage), and wind farms (bird strike). However, further research will likely be needed to enhance the efficacy. This study has identified a new principle for repelling birds through physiological methods using flickering light stimuli. In the current challenging situation of mitigating human-avian conflict, the impact of this discovery should not be underestimated.

## Acknowledgements

The authors would like to thank Kaori Muramatsu for her valuable assistance in organizing the data and supporting the research. During the preparation of this work, the author utilized GPT-4.0 to improve spelling and grammar.

## Authorship contributions

The sole author, Takeshi Honda is responsible for all aspects of this study.

## Data availability

The data will be provided upon reasonable request to the corresponding author.

## Statements and Declarations Ethical Approval

All applicable international, national, and institutional guidelines for the care and use of animals were followed. This animal study was approved by the Yamanashi Prefectural Research Assessment Committee (#050901).

