## Supplementary figures and images for "The physiology of deterrence: Flicker vertigo and its application in avian management"

### appendix A

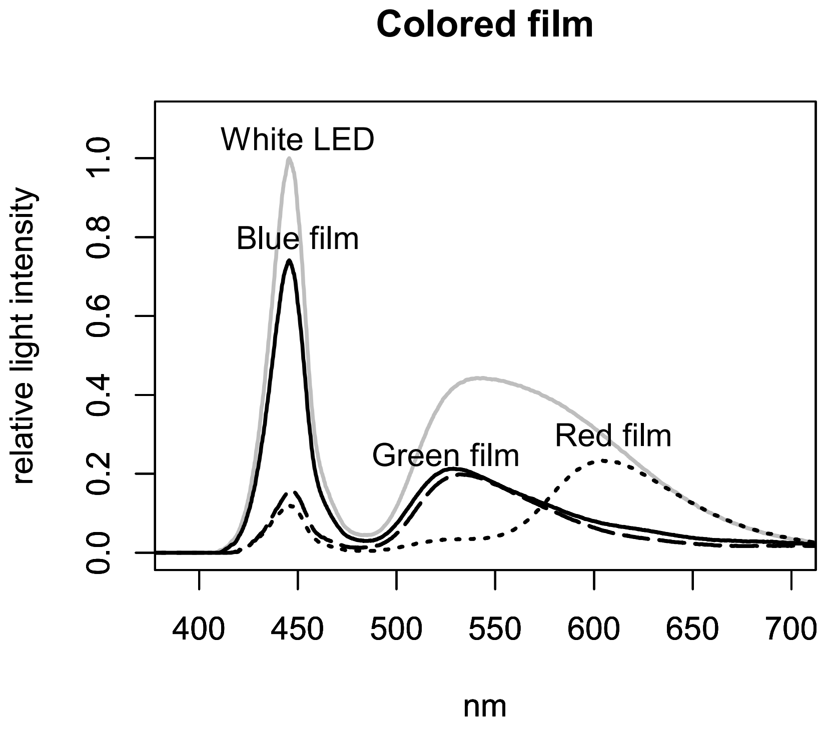

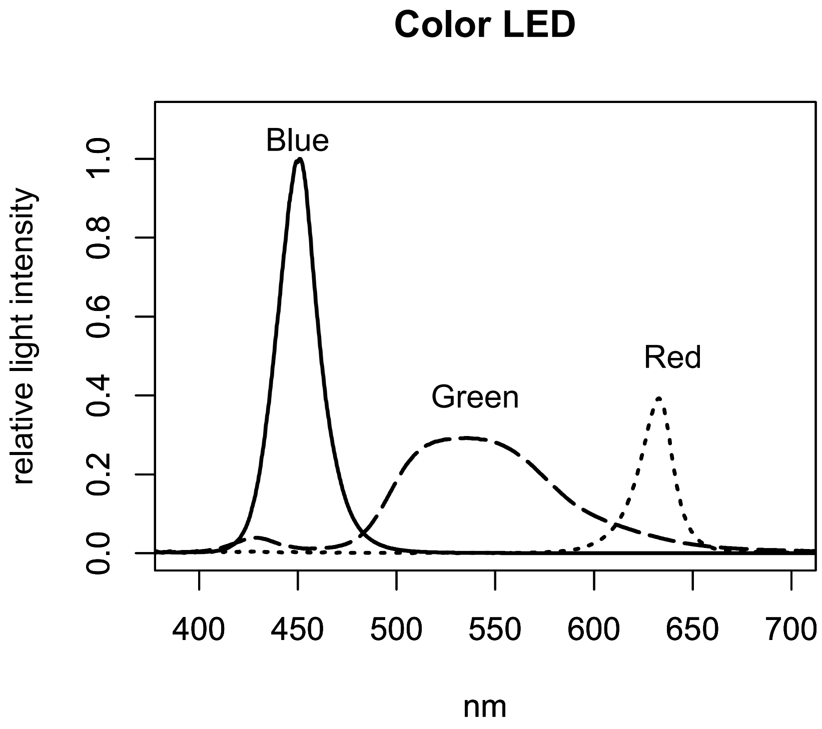
